# New distribution records of *Aedes aegypti*, *Aedes mediovittatus*, and *Toxorhynchites portoricensis* (Diptera: Culicidae) in Puerto Rico and their relevance to Integrated Vector Management

**DOI:** 10.1101/2025.10.22.683737

**Authors:** Jun Soo Bae, Telmah Telmadarrehei, Sangwoo Seok, Lianmarie Soto Jiménez, Amaury Morales González, Luis F. Quintanilla Vásquez, Valerie T. Nguyen, Riley Young, Raymond Gellner, Lawrence E. Reeves, Joanelis Medina, Grayson Brown, Yoosook Lee

## Abstract

As of October 2025, Puerto Rico has been experiencing an ongoing dengue outbreak that started in March 2024. The latest island-wide mosquito survey conducted in Puerto Rico during 2018– 2019 covered 41 of the 78 municipalities and detected the presence of *Aedes aegypti* in 27 of the municipalities. Given the prolonged elevated circulation of dengue virus on the island, we carried out an additional *Ae. aegypti* survey in June 2025 across 44 out of 78 municipalities, including areas with no prior records of the species. Here, we report the occurrence of *Ae. aegypti* in 43 out of 44 municipalities surveyed, including ten new municipalities where *Ae. aegypti* has not been reported. These findings provide insight into the dispersal of *Ae. aegypti* in Puerto Rico, indicating that the species is much more widespread than previously known. We also provide the expanded species occurrence of *Aedes mediovittatus* and *Toxorhynchites portoricensis*, which share the same larval habitat as *Ae. aegypti*. Notably, 96% of cemeteries surveyed across 24 municipalities served as oviposition sites for *Ae. aegypti*. However, a few cemeteries demonstrated effective preventative practices that minimize mosquito breeding. This effective management, which resulted in no mosquito breeding in some cemeteries and a tire shop, brings hope that communication and education of property/facility managers can reduce the mosquito populations.

## INTRODUCTION

Accurate occurrence records of mosquito species are important for mosquito management and pathogen surveillance. Regular surveillance of mosquito species can enable monitoring of their distribution and abundance, allowing public health agencies to respond promptly to potential risks (Bakhiyi et al. 2024). Such data is also critical for evaluating the effectiveness of control programs and for developing future management strategies. Moreover, because environmental factors such as urbanization and climate change can increase vector abundance and risks of infection (Ryan et al. 2019, Wilke et al. 2021), regular surveillance is a key component of mosquito management strategies.

As of October 2025, Puerto Rico has been experiencing an ongoing dengue outbreak that was declared in March 2024 (Ware-Gilmore et al. 2025). This region is currently experiencing the highest dengue case load in the United States (Figure 1). Documented outbreaks in Puerto Rico have involved the co-circulation of multiple dengue serotypes (1-4) which exacerbates the health risks to residents (Rigau-Pérez et al. 2002, Sharp et al. 2019, Rodriguez et al. 2024). High circulation of dengue virus in Puerto Rico can also spill over to other neighboring regions including the continental United States leading to high travel-related dengue cases as well as increasing the risk of local dengue transmission in neighboring states like Florida (Taylor-Salmon et al. 2024). Therefore, studying dengue virus and its vector species in Puerto Rico is not only beneficial for the residents in Puerto Rico but also for population in the neighboring areas and the whole nation.

**Figure 1.**
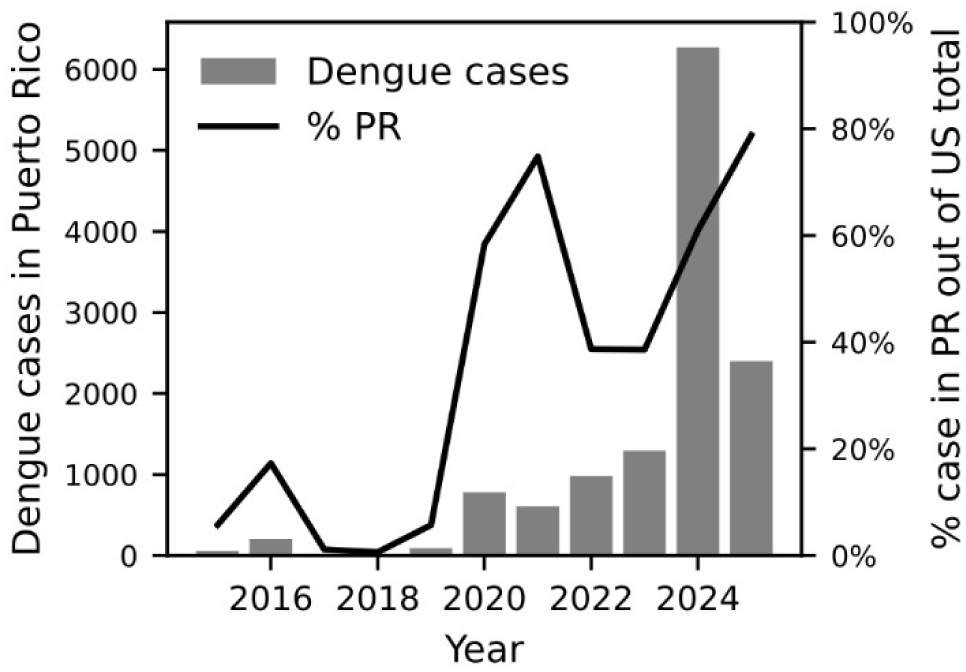
Dengue cases in Puerto Rico between 2015-2025 (gray bar) and % cases corresponding to Puerto Rico compared to the national case number (black line). Data is from CDC (CDC 2025)

Invasive species *Aedes aegypti* serve as vectors for dengue virus in Puerto Rico while the native species *Aedes mediovittatus* has shown high competency for transmission (Tomashek et al. 2009, Poole-Smith et al. 2015). *Aedes albopictus*, another invasive mosquito species that acts as a vector of dengue virus, was thought to be introduced to Puerto Rico before 2005 (Cook et al. 2006). However, *Ae. albopictus* was not observed in the multiple island-wide mosquito species surveys in following years (Barrera et al. 2011, 2012, Yee et al. 2021). The most recent island-wide survey encompassed 41 municipalities in Puerto Rico documenting the presence of *Ae. aegypti* from 27 municipalities and *Ae. mediovittatus* in 19 municipalities (Yee et al. 2021). Fourteen of the 41 municipalities (34%) harbored both *Ae. aegypti* and *Ae. mediovittatus*. In Utuado, Puerto Rico, *Ae. aegypti* was also found co-occurring with *Tx. portoricensis* (Yee et al. 2021).

Recent study investigated presence of mosquito larvae in over 9000 water-holding containers from 16 cemeteries across six municipalities (Caguas, Humacao, Juncos, Las Piedras, Naguabo, and Yabucoa) in the East region of Puerto Rico between 2019 and 2020 (Otero et al. 2022). *Aedes aegypti* and *Ae. mediovittatus* were identified as the most abundant species. The two species accounted for 84.9% of all collected immature mosquitoes in all six municipalities surveyed. *Aedes aegypti* was present in every cemetery except for Municipal Ramon Delgado (Juncos) and Vale de Paz (Las Piedras). *Aedes mediovittatus* was identified in every cemetery except Borinquen Memorial I (Caguas), Valle de Paz (Las Piedras) and La Inmaculada (Juncos). However, *Tx. portoricensis* was not reported in the study.

During our investigation checking prior species occurrence records of *Ae. aegypti* in Puerto Rico using Global biodiversity Information Facility (GBIF) and other publications, we discovered that 20 municipalities out of total 78 municipalities in Puerto Rico still had no records of the presence of *Ae. aegypti* (Yee et al. 2021, Otero et al. 2022, GBIF Secretariat 2023). Given the ongoing dengue outbreak in Puerto Rico, we are compelled to report updated species occurrence records so that municipalities without *Ae. aegypti* records can limit mosquito reproduction.

This study also presents updated species records for *Ae. mediovittatus* and *Tx. portoricensis,* which share some oviposition sites with *Ae. aegypti* in Puerto Rico (Otero et al. 2022). We also provide examples of good practices that can reduce mosquito breeding in cemeteries and tire shops based on the three locations where we failed to collect any mosquito larvae. This information can be further developed into educational materials for the public for source reduction efforts, which is an integral part of integrated vector management.

## MATERIALS AND METHODS

### Mosquito collection

We collected both adult and immature stages (larvae and pupae) of mosquitoes from 44 municipalities (56%) in Puerto Rico covering 100 locations in June of 2025 during the wet season (Table S1). We set up BG-Sentinel (BG-S) traps (Biogents AG, Regenburg, Germany) in residential areas with permission from homeowners. An average of 1-2 BG-S traps were set per household. Traps were set to run for 24-36 hours. Mosquitoes in BG-S collection bags were brought to a laboratory and kept in -20°C until species identification by morphological examination.

Flower vases in cemeteries (Figure 2A and 2B), discarded tires on the roadside (Figure 2C), and other containers holding water (Figure 2D) were examined for the presence of immature stages (larvae and pupae). We examined multiple locations within a cemetery for mosquito larvae and samples were collected from one to six locations within each cemetery. They were extracted using turkey basters, a ladle, and/or plastic transfer pipettes. The larvae and pupae were held in Whirl-Pak plastic bags (Whirl-Pak, Austin, TX) for transport between collection sites and the base of operation, which was either Puerto Rico Vector Control Unit (PRVCU) laboratory in San Juan or Ponce. GPS coordinates of each trap and larval site were recorded using Google Maps app on mobile devices. For larvae sampled from multiple vases within two-meter radius were stored in one bag with one GPS coordinate. Larvae were put into containers separated by coordinates and left until pupation. Pupae were transferred into 7-dram vials with water for emergence. The eclosed adults were frozen in -20°C freezer before species identification. All samples were preserved in 70% ethanol for DNA extraction.

**Figure 2.**
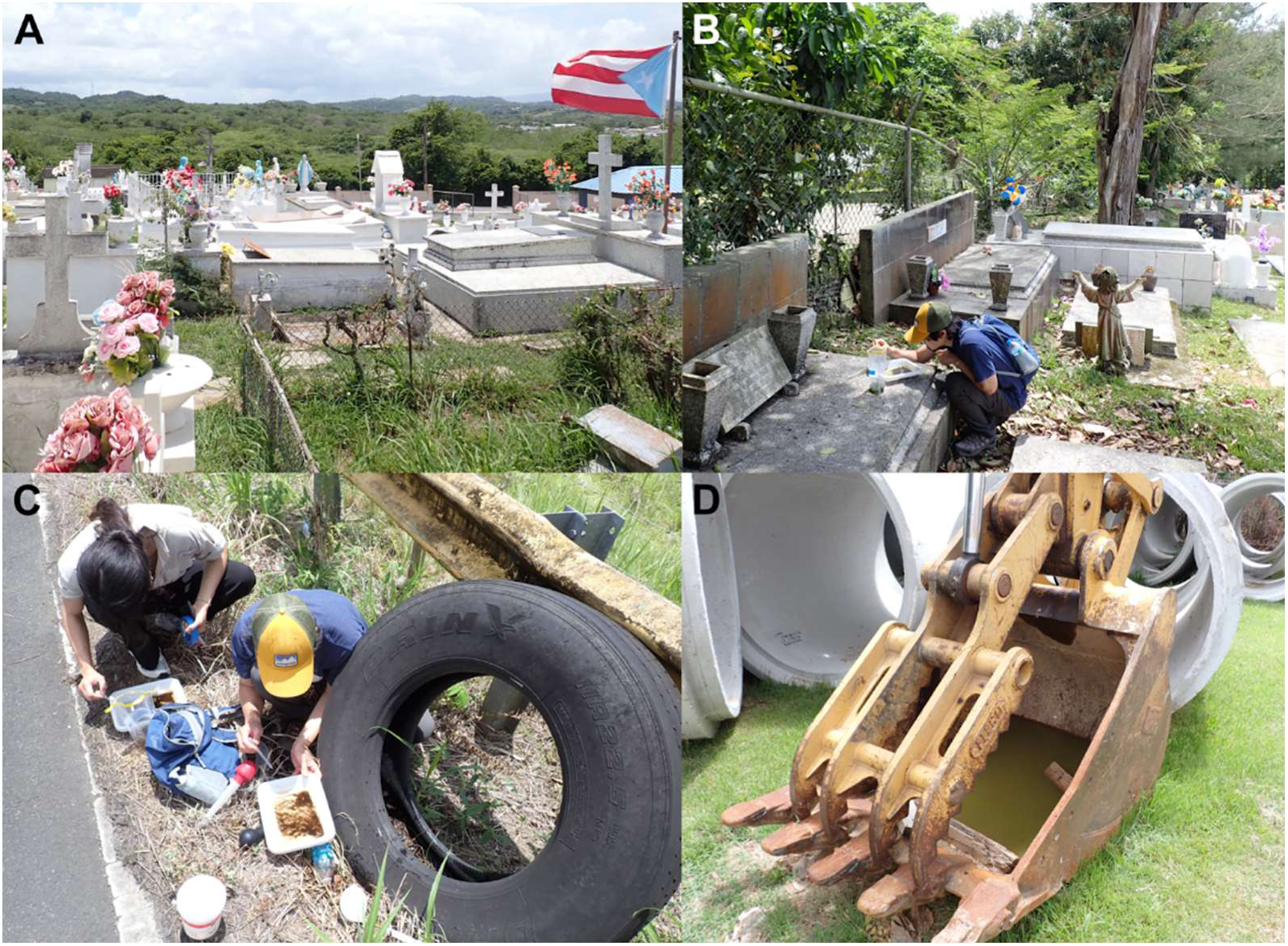
Example habitat photos. A: a cemetery with flower vases. B: a cemetery with flower vases and more shade. C: discarded tire holding water. D: excavator bucket holding water.

### Species identification

Species identification of adult mosquitoes was determined by morphology using the in-house identification keys provided by the PRVCU. Concurrently, IDX device (Vectech, Baltimore, MD), machine learning-based *Aedes aegypti* identification version 6.0.0, was used to separate *Ae. aegypti* from non-target species more efficiently. This device has been used in other studies to identify *Ae. aegypti* and showed 88.8% accuracy at version 4.0 (Gupta et al. 2024). We accepted *Ae. aegypti* species calls if the confidence level is above 80%. The samples with less confident (<80%) or unknown calls were further examined for morphological features to confirm its species according to PRVCU identification keys. Samples we could not confidently confirm were stored in 70% alcohol for molecular species identification.

Larvae, pupae, and damaged adult specimens were identified to species using multiple molecular assays. First, samples were checked if they were *Ae. aegypti* by an internal transcribed spacer 2 (ITS2) PCR assay developed by Menegon *et al*. (2025) with modified set of primers using Aedes-F2 (5’-AGG ACA CAT GAA CAC CGA CA-3’), JAP-R (5’-TAT ACT ACG CTG CCG AGA GG-3’), and AEG-R2 (5’-TGA GTG AAT GAT GGA ATA CAA CA-5’) primers. *Aedes albopictus* primers of Menegon *et al*. (2025) were not included because they repeatedly failed to amplify for positive control. A 25 µL of PCR mixture was prepared for each sample to contain 1 µL of extracted DNA template, 12.5 µL of 2X OneTaq Master mix with buffer (Life Technologies, Carlsbad, CA), 0.05 µL each primer in 10 µM (final concentration 10 pmol per primer), and 11.3 µL of PCR-grade water to amplify ITS2 region. DNA was initially denatured at 94 °C for 5 min, followed by 40 cycles of denaturation at 94 °C for 30s, annealing at 50.2°C for 30s, and extension at 72 °C for 60s. Then a final extension step of 72 °C was set for 5 min before the PCR products were held at 4 °C before storage at -20 °C. The PCR products were stained with SYBR^TM^ Safe DNA Gel Stain (Invitrogen, Waltham, MA) and separated by its fragment size by electrophoresis on a 1.5% agarose gel.

For samples with no amplification on *Aedes* ITS2 PCR assay (Menegon et al. 2025), we amplified cytochrome c oxidase I (COI) sequence using DNA Barcoding primers LCO1490 (5’-GGT CAA CAA ATC ATA AAG ATA TTG G-3’) and HCO2198 (5’-TAA ACT TCA GGG TGA CCA AAA AAT CA-3’) (Hebert et al. 2003). PCR protocol followed the method described in Reeves et al. (2021). The PCR product was sent to Eurofins for Sanger Sequencing. The resulting DNA sequences were entered into Basic Local Alignment Search Tool - nucleotide (BLASTn) to find a matching species (Altschul et al. 1990).

### Data Analysis

Maps of sampling locations and species occurrence records were generated using QGIS version 3.40.4 (QGIS 2025). The shapefile of Puerto Rico municipalities was obtained from Datos.PR (Datos.PR 2018). For data reporting purposes, we used six regions – namely Metro, East, North, Central, South, and West regions - commonly referred to in governing and tourism (Rivera 2025) for aggregating municipality data. We used GBIF (GBIF Secretariat 2023), Yee et al. (2021), and Otero et al. (2022) records to determine which municipality records are new in Puerto Rico. We also used the six United States Geological Survey climatic subdivisions in Puerto Rico (USGS CFWSC 2016) to examine any trends in association with climatic conditions and mosquito species distribution.

## RESULTS

### *Aedes aegypti* occurrence in Puerto Rico

The survey conducted in June 2025 across 44 municipalities over two weeks revealed that adult or immature stage (larvae and pupae) of *Ae. aegypti* were present in 43 municipalities (Figure 3). Our survey includes 10 new municipality records of *Ae. aegypti* occurrence (marked with asterisk (*) in Table 1). We did not collect any mosquitoes from Maricao municipality in the West region.

**Figure 3.**
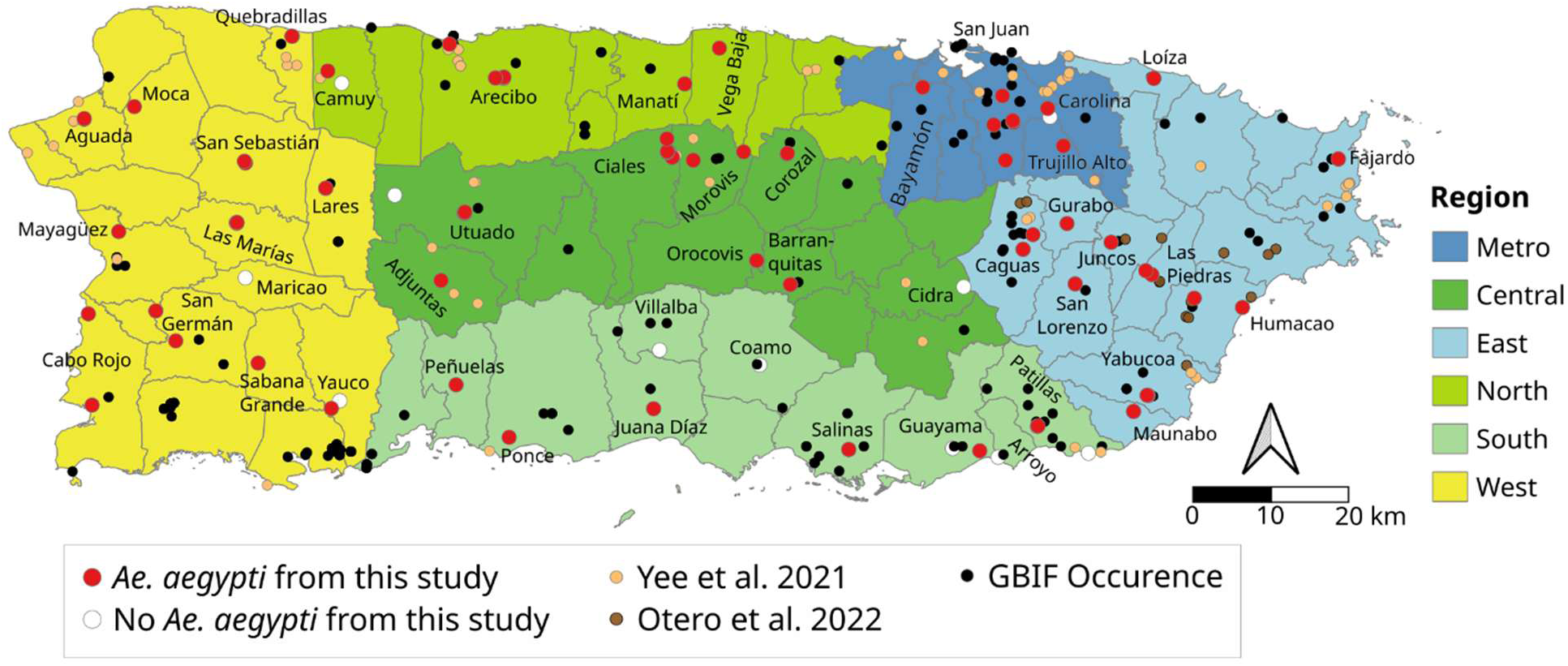
Occurrence records of *Ae. aegypti*. Metro region is marked in blue, East region in light blue, North region in green, Central region in dark green, South region in lime green, and West region in yellow background color. Black dots indicate GBIF occurrence record as of June 2025, orange dots indicate occurrence records from Yee et al. (2021), brown dots indicate occurrence records from Otero et al. (2022), and red dots indicate data from this study.

**Table 1.**
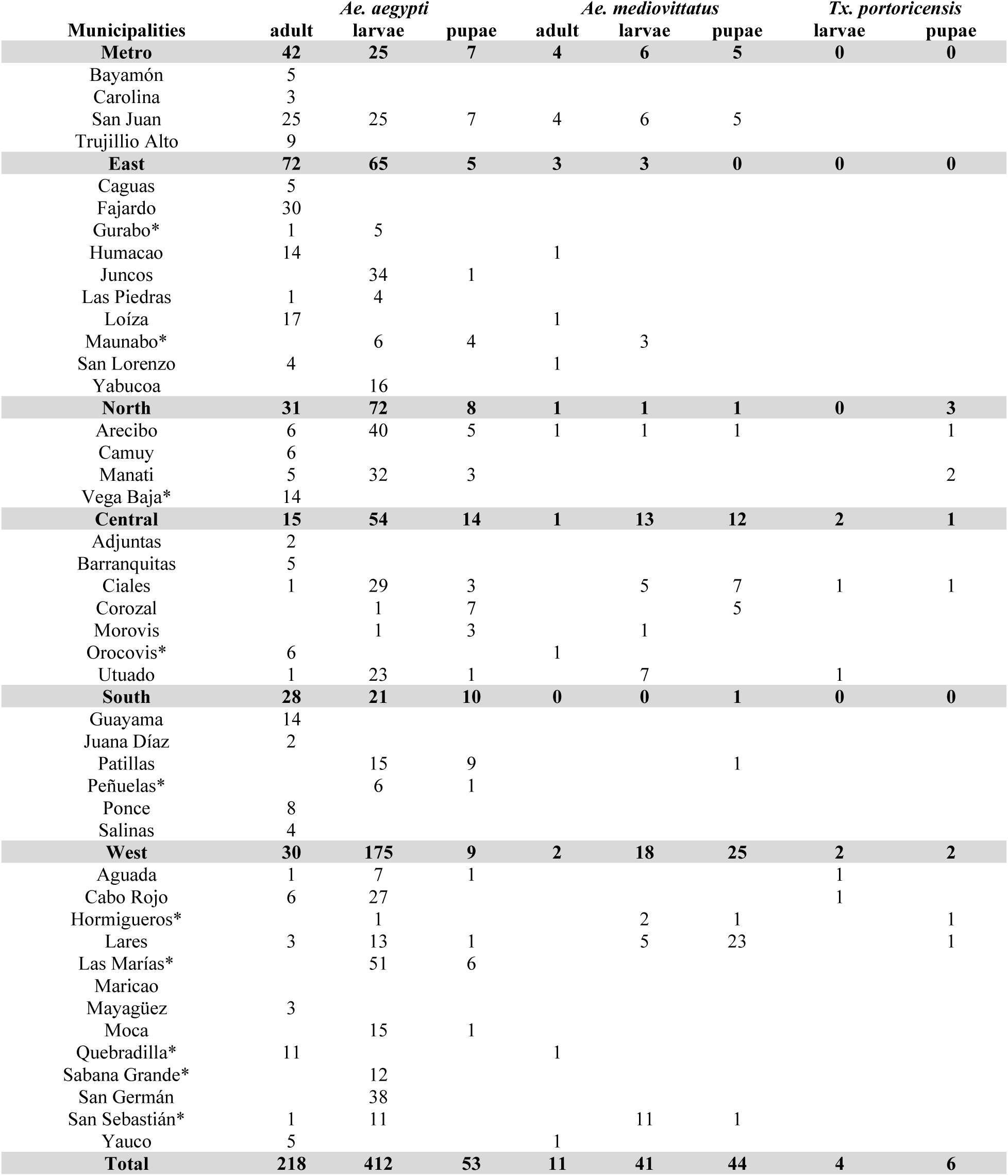
Collection records of *Ae. aegypti, Ae. albopictus, Ae. mediovittatus,* and *Tx. portoricensis* in municipalities of Puerto Rico during June 2025. The adult and larvae/pupae columns indicate the life stage at time of collection in the field. Asterisk (*) indicates new municipality record of *Ae. aegypti* occurrence. The summary table is provided here for brevity and detailed sample metadata are provided in supplemental Table S1.

The majority of larval sites we examined were in cemeteries (52/63 =82.5%). Collections from water held in tires (n=6 in 4 municipalities) were serendipitous encounters as we travel between municipalities and thus occupy relatively small portions of our larval collection. *Aedes aegypti* were found in 81.0% of the larval collection sites.

### *Aedes albopictus* occurrence in Puerto Rico

We did not collect any *Ae. albopictus* during our mosquito survey.

### *Aedes mediovittatus* occurrence in Puerto Rico

*Aedes mediovittatus* was found in 26 locations in 17 municipalities (Figure 4). Ten of the 17 municipalities (Ciales, Corozal, Hormigueros, Lares, Loíza, Orocovis, Quebradillas, San Lorenzo, San Sebatian, and Utuado) are new municipality records (Yee et al. 2021, Otero et al. 2022, GBIF Secretariat 2023). This species was most commonly found in West region (8/23 = 34.8%) and least common in South region (1/9 = 11.1%; Table 2). With respect to USGS Climatic subdivision (USGS CFWSC 2016), *Ae. mediovittatus* was most common in the North Coastal (6/17=35.3%) and Western Interior (11/34 = 32.4%) region. Catch numbers using BG-S traps were generally low (n=0-4 per trap). *Aedes mediovittatus* are often found together with *Ae. aegypti* (23/26 =88.5%). In comparison, cooccurrence of *Ae. mediovittatus* and *Ae. aegypti* were previously noted at 77% (10/13, Otero et al. 2022) and 1.76 % (13/74, Yee et al. 2021).

**Figure 4.**
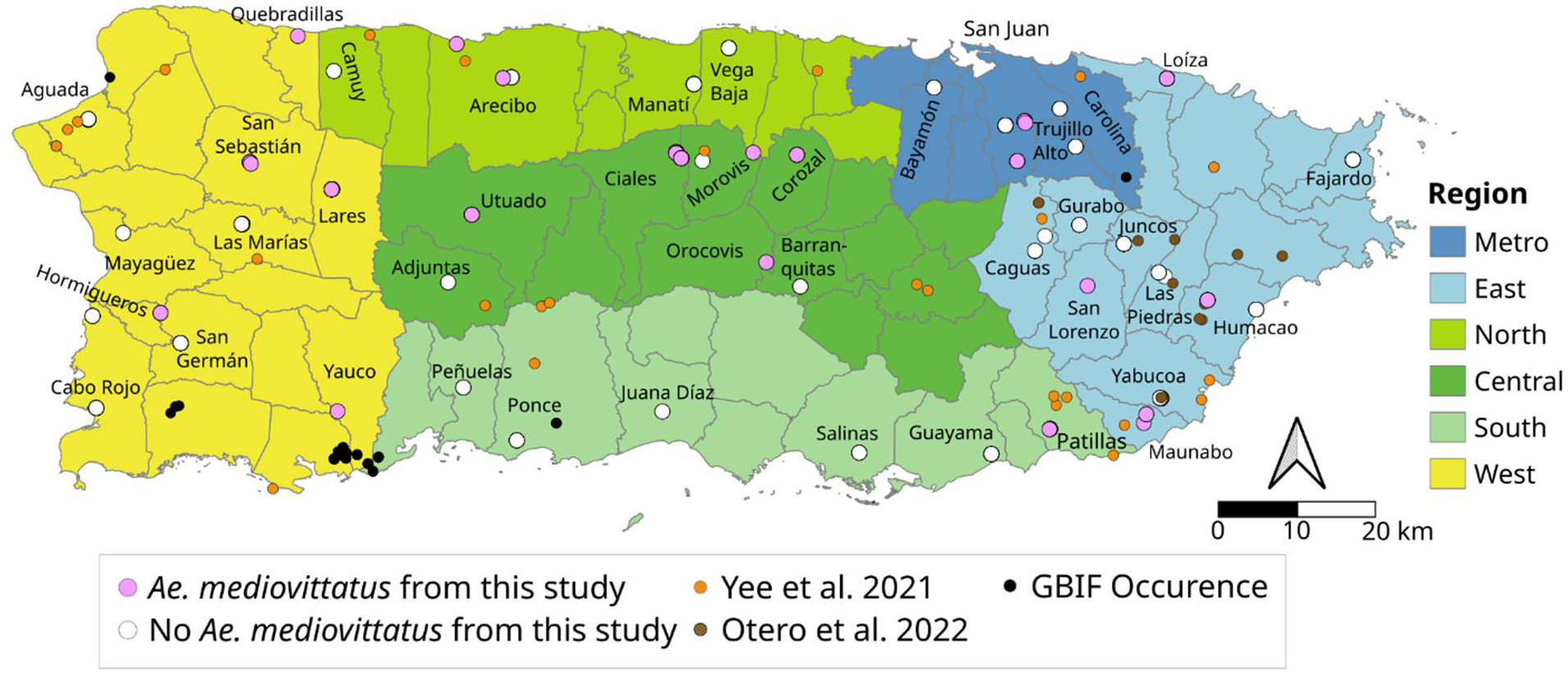
Occurrence records of *Ae. mediovittatus.* Metro region is marked in blue, East region in light blue, North region in green, Central region in dark green, South region in light green, and West region in yellow background color. Black dots indicate GBIF occurrence record as of June 2025, Orange dots indicate occurrence records from Yee et al. (2021), brown dots indicate occurrence records from Otero et al. (2022), and pink dots indicate data from this study.

**Table 2.** The number of locations where *Ae. mediovittatus* was detected grouped by Region or USGS Climatic subdivision (2016).

| Region | Total collection locations | *Ae mediovittatus* positive locations | % positive | USGS Climatic subdivision (2016) | Total collection locations | *Ae mediovittatus* positive locations | % positive |
| --- | --- | --- | --- | --- | --- | --- | --- |
| Metro | 15 | 4 | 26.7% | North Coastal | 17 | 6 | 35.3% |
| East | 21 | 5 | 23.8% | Eastern Interior | 10 | 1 | 10.0% |
| North | 10 | 2 | 20.0% | Northern Slopes | 15 | 2 | 13.3% |
| Central | 22 | 6 | 27.3% | Western Interior | 34 | 11 | 32.4% |
| South | 9 | 1 | 11.1% | South Coastal | 4 | 1 | 25.0% |
| West | 23 | 8 | 34.8% | Southern Slopes | 20 | 5 | 25.0% |

### Toxorhynchites portoricensis occurrence in Puerto Rico

*Toxorhynchites portoricensis* was found in nine locations in eight municipalities from our study (Figure 5). These include Ciales and Utuado in the Central region, Arecibo and Manatí in the North region, and Aguada, Cabo Rojo, Lares, and Las Marias in the West region. Seven of the eight municipalities (Aguada, Arecibo, Cabo Rojo, Ciales, Lares, Hormigueros, and Manatí) are new municipality records for this species. Previously, occurrence of this species was noted in Bayamón, Cayey, Isabela, Luquillo, Patillas, Río Grande, San Juan, and Utuado municipalities (Yee et al. 2021, GBIF Secretariat 2023). Collectively, this species has been found in all regions except the South region (Figure 5, Table 3). With respect to USGS Climatic subdivision (USGS CFWSC 2016), this species was most common in Northern Slopes (13.3%) followed by Western Interior (11.8%) and Southern Slopes (10.0%). We did not encounter this species in the Eastern Interior or South Coastal region.

**Figure 5.**
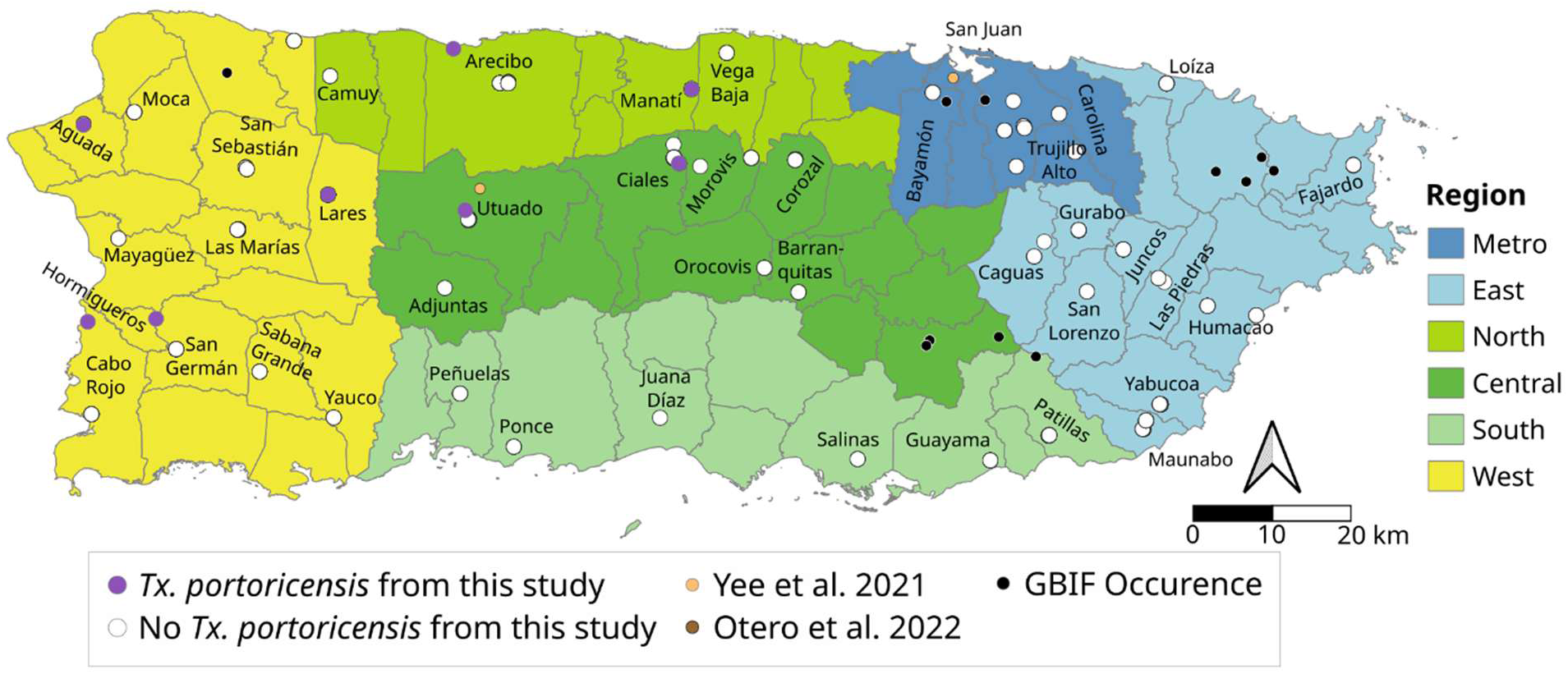
Occurrence records of *Tx. portoricensis.* Metro region is marked in blue, East region in light blue, North region in green, Central region in dark green, South region in light green, and West region in yellow background color. Black dots indicate GBIF occurrence record as of June 2025, blue dots indicate occurrence records from Yee et al. (2021), and purple dots indicate data from this study. *Toxorhynchites* data is not recorded in Otero et al. 2022.

**Table 3.** The number of locations where *Tx. portoricensis* was detected grouped by Region or USGS Climatic subdivision (2016).

| Region | Total collection locations | *Tx portoricensis* positive locations | % positive | USGS Climatic subdivision (2016) | Total collection locations | *Tx portoricensis* positive locations | % positive |
| --- | --- | --- | --- | --- | --- | --- | --- |
| Metro | 15 | 0 | 0% | North Coastal | 17 | 1 | 5.9% |
| East | 21 | 0 | 0% | Eastern Interior | 10 | 0 | 0% |
| North | 10 | 2 | 20.0% | Northern Slopes | 15 | 2 | 13.3% |
| Central | 22 | 3 | 13.6% | Western Interior | 34 | 4 | 11.8% |
| South | 9 | 0 | 0% | South Coastal | 4 | 0 | 0% |
| West | 23 | 4 | 17.4% | Southern Slopes | 20 | 2 | 10.0% |

### Successful cases of integrated vector management in Puerto Rico

There were three locations where we did not detect any mosquito presence during our collection. Two were cemeteries in Maricao and Morovis municipalities and one tire shop in Morovis municipality. The entrance of the Maricao cemetery had multiple clear signs banning the use of flower vases in the cemetery (Figure 6A). Conversation with the Maricao cemetery manager revealed that he used to collaborate with PRVCU in the past and was aware of likely places where mosquitoes can breed and actively managed to minimize creating pools of water in the cemetery. The common features of the Mauricao and Morovis cemeteries included vases filled to the brim with sand or dirt, vases with drainage holes, and the practice of turning vases upside down when not in use (Figure 6D-F). Prevention at a tire shop included preventing buildup of water by shipping out used tires weekly and adding holes into a shop sign made from a used tire (Figure 6B and 6C).

**Figure 6.**
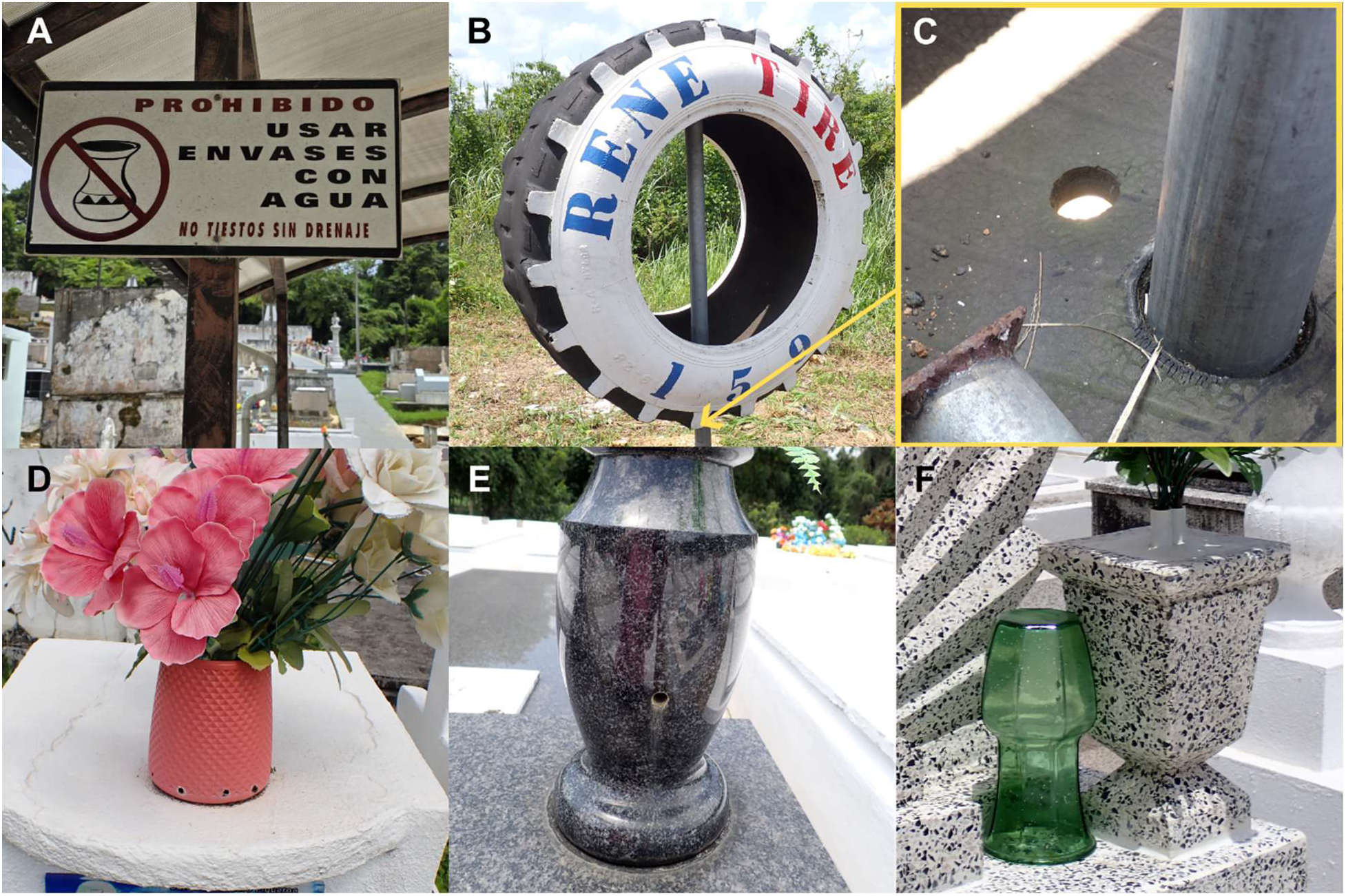
Examples of management practices from locations without any mosquitoes. A: Sign displayed at the entrance of the cemetery in Maricao prohibiting the use of containers without drainage that can hold water. B: Tire landmark C: with hole drilled on the base of the inside of the tire for drainage. D: Flower vase with holes on the base for drainage. E: Stone flower vase with drainage provided by the cemetery. F: Use of faux flowers with pot cemented and vase left upside down to prevent accumulation of stagnant water.

## DISCUSSION

### Distribution of dengue vector in Puerto Rico

During the ongoing dengue outbreak in Puerto Rico, we were able to observe *Ae. aegypti* from 43 municipalities out of 44 total municipalities surveyed in June 2025. Our records include 10 new municipality records that have not previously reported the presence of *Ae. aegypti*. The ubiquitous occurrence of *Ae. aegypti* indicates that this species does not have limiting environmental conditions within Puerto Rico. The collecting period of this study was the early stage of the rainy season, and we anticipate that more locations will become available for *Ae. aegypti* reproduction in the later period of the year around late September and early October based on the past trends in dengue cases (Salud PR 2025).

From our survey of 114 locations in 43 municipalities, we did not find *Ae. albopictus*. The presence of *Ae. albopictus* in Puerto Rico was first reported in 2006 (Cook et al. 2006). Since this initial report, the subsequent extensive surveys have not found *Ae. albopictus* on the main island of Puerto Rico (Barrera et al. 2011, 2012, Yee et al. 2021). In the southernmost municipalities of the West region, GBIF records with concrete dates of observation reports *Ae. albopictus* between 2016-2020 from Lajas and Guánica municipalities (GBIF Secretariat 2023). However, as these two regions were not included in this study, we were unable to verify the records. Therefore, the presence of *Ae. albopictus* in Puerto Rico needs further investigations to verify.

### Other mosquitoes sharing oviposition sites with *Ae. aegypti*

Based on available information, *Aedes mediovittatus* is a native mosquito species in Puerto Rico and shares oviposition sites with *Ae. aegypti* (Otero et al. 2022)*. Aedes aegypti* has been frequently reported to outcompete other mosquito species in interspecific interactions (Santana-Martínez et al. 2017, Lushasi et al. 2024). Nevertheless, *Ae. mediovittatus* still coexists with *Ae. aegypti* in shared habitats, possibly because *Ae. mediovittatus* exhibits a competitive advantage over *Ae. aegypti* (Yee et al. 2025).

*Aedes mediovittatus* are continuously monitored for dengue infection by PRVCU because they are often collected with *Ae. aegypti.* In 2025, most of the dengue-positive pools that PRVCU tested were from *Ae. aegypti.* This indicates that *Ae. mediovittatus* could be more resilient in dengue virus infection compared to *Ae. aegypti* and thus a less competent vector for dengue transmission. While *Ae. aegypti* has many studies investigating its immune responses against dengue virus infection with good genomic resources (Behura et al. 2011, Castillo-Méndez and Valverde-Garduño 2020), little is known about *Ae. mediovittatus.* There are very limited genetic resources for *Ae. mediovittatus* with only two COI sequences available as of September 2025 (Gibson et al. 2012). While it will take a long time to build equivalent genetic resources for *Ae. mediovittatus* for comparative genomics/transcriptomics studies, the comparison of immune responses against dengue infection between *Ae. mediovittatus* and *Ae. aegypti* could illuminate mechanisms of virus replication in these mosquito species. The understanding of virus transmission in two different mosquitoes with varying degrees of vector competency can be utilized to block virus transmission in mosquitoes in the future.

*Toxorhynchites portoricensis* is the only *Toxorhynchites* genus known to occur in Puerto Rico. This species belongs to the *Lynchiella* subgenus. In this study, we provided a broader distribution of *Tx. portoricensis* occurrence records than previously known with seven new municipality records. Limited observations of this species in Puerto Rico were mostly away from coastal areas (Yee et al. 2021, GBIF Secretariat 2023). However, this species appears to be more widespread than previously thought, including locations close to the coast in Aguada, Arecibo, and Cabo Rojo municipalities (Figure 4). *Toxorhynchites* species are typically found in locations like discarded tires and tree holes (Schreiber 2007). We found *Tx. portoricensis* in mostly tires, however, also found them in flower vases in Ciales and in water collected in an excavator scoop in the residential area in Cabo Rojo (Figure 2D).

During our field work, we observed *Tx. portoricensis* larvae consuming their own species in the same water. The carnivorous larva of this genus consumes other mosquito larvae and has long been considered for biological control options as part of the integrated vector management (Focks 2007, Albeny-Simões et al. 2011, Seok et al. 2022). *Aedes aegypti* in particular are attracted to the bacterial makeup of waters already predated by *Toxorhynchites theobaldi*, making *Toxorhynchites* a more beneficial contender as biocontrol agent (Albeny-Simões et al. 2014). One of the characteristic behaviors in some *Toxorhynchites* species is the prepupal killing of surrounding larvae without consuming them before becoming pupae (Focks 2007). While this behavior has not been observed in this species yet, it might represent another possible way by which it suppresses other mosquitoes.

The colonizability of *Tx. portoricensis* is unknown. For this species to be considered for a biocontrol agent, one must be successfully colonize and reared in large numbers in laboratory conditions. One of the co-authors (A. M. González) attempted a single pair mating in a typical cage (1 cu. ft. in size) condition in the past, but the female did not lay any eggs. The spermatheca of the female was not examined to check if the female carried sperm after the mating attempt. More studies are needed such as trying larger cages, increasing the numbers of adults available in a cage for mating, and so on before we can rule out the feasibility of establishing a colony of *Tx. portoricensis*.

### Integrated vector management to reduce *Ae. aegypti*

Every municipality has at least one cemetery in Puerto Rico, often located in one of the major towns in the municipality adjoining the large residential area. It is a permanent fixture in their landscape and often has dedicated maintenance staff managing the ground. The successful mosquito management cases, such as those seen in Mauricao and Morovis cemeteries and a tire shop, highlight that the right focus on public education campaigns can have a long-lasting impact on reducing mosquito breeding and can reduce the risk of mosquito biting for large swaths of the population centers. Focused education on the maintenance and property management personnel on the permanent infrastructure like cemeteries could have lasting impact in reducing the mosquito reproduction in Puerto Rico.

## ACKNOWLEDGEMENTS

We thank PRVCU affiliates, Reynaldo Morales, Christian Sánchez, Noemí Martínez, Marla García, Amaury Morales, Rafael Saavedra, Mariela Charlotten, Paola Colón, Noel J. Del Pilar, Luis Doel Santiago, Warren Ortiz, Luis R. Perez, Javier A. Cordova, Herick Leon, Hernan Olivera, Neysha Rosado Ramos, Noraliz Rodríguez Burgos, Nexilianne Borrero-Segarra, Paola Quiñones Cruz, and Joel Munoz Flores for assisting with mosquito collection using BG traps in residential areas. We thank Noemí Martínez at PRVCU for arranging with community leaders in identifying potential BG trap sites in Western and Southern Puerto Rico locations.

## Disclaimer

The findings and conclusions in this manuscript are those of the authors and do not necessarily represent the official position of the National Institute of Health, the Southern Integrated Pest Management Center, and USDA.

## Supplemental Materials

**Table S1.** Detailed collection records of *Ae. aegypti, Ae. mediovittatus,* and *Tx. portoricensis* in Puerto Rico during June 2025. The adult and larvae/pupae columns indicate the life stage at time of capture. Multiple trap types were used in some municipalities.

**Table S2**. *Aedes aegypti* occurrence record comparisons

